# ErbB inhibition impairs cognition via disrupting myelination and aerobic glycolysis in oligodendrocytes

**DOI:** 10.1101/2023.01.03.522525

**Authors:** Xu Hu, Qingyu Zhu, Tianjie Lou, Qianqian Hu, Huashun Li, Xiaojie Niu, Li He, Hao Huang, Yijia Xu, Mengsheng Qiu, Ying Shen, Jie-Min Jia, Yanmei Tao

## Abstract

White matter abnormalities are an emerging feature of schizophrenia, yet the underlying pathophysiological mechanisms are largely unknown. Disruption of ErbB signaling that is essential for peripheral myelination has been genetically associated with schizophrenia and white matter lesions in schizophrenic patients. However, the roles of ErbB signaling in oligodendrocytes remain elusive. Here, we used a pan-ErbB inhibition strategy and demonstrated the synergistic functions of endogenous ErbB receptors in oligodendrocytes. Through analyses of the cellular, histological, biochemical, behavioral, and electrophysiological differences in mice with manipulation of ErbB activities in oligodendrocytes at different differentiation stages, we found that ErbB signaling regulates myelination and aerobic glycolysis in oligodendrocytes, and both functions are required for working memory. ErbB inhibition in oligodendrocytes at early differentiation stages induces hypomyelination by suppressing the differentiation of newly-formed oligodendrocytes. In contrast, ErbB inhibition in mature oligodendrocytes alters neither myelination nor oligodendrocyte numbers, but accelerates axonal conduction decline under energy stress. Mechanistically, mature oligodendrocytes with ErbB inhibition reduce the expression of lactate dehydrogenase A, failing to provide lactate to electrically active axons. Supplementation of L-lactate restores axonal conduction and working memory capacity that are suppressed by ErbB inhibition in mature oligodendrocytes. These findings reveal the indispensable roles of ErbB signaling in white matter integrity and function, and provide insights into the multifaceted contributions of white matter abnormalities to cognitive impairment.

## Introduction

Adolescence is the critical period for the central nervous system (CNS) to completely develop and mature. In particular, CNS myelin generated by oligodendrocytes (OLs) is one of the most developmentally active component in the adolescent brain. This may lead to CNS myelin being a highly susceptible target in psychiatric disorders such as schizophrenia that typically develops during adolescence ^1–3^. A growing body of literature points to abnormalities in the structure, component proteins, or regulating molecules of CNS myelin in schizophrenic patients ^1, 4–8^. Particularly, white-matter microstructural changes that can be examined by structural brain imaging techniques are sensitive to the patient age, symptom characteristics, and genetic loading ^9–11^. Given that it is one of the most promising features whose progression can be examined periodically in patients, understanding the biological mechanisms for schizophrenia-related white matter abnormalities is crucial for the development of diagnostic criteria and therapeutic targets.

A few myelin protein encoding genes, including *PLP1, MOG, MAG*, and *MPZL1*, exhibit genetic association with schizophrenia ^12–16^. However, most of these variants are only revealed in Han Chinease popuation ^17^. In contrast, tyrosine kinase receptor ERBB4 and its ligand neuregulin 1 (NRG1), which has been proved to be essential for peripheral nervous system (PNS) myelination ^18^, exhibit aberrant expression and genetic polymorphisms that are associated with schizophrenia in many populations ^17, 19, 20^. Studies combining genetic linkage analysis and brain imaging techniques have associated *NRG1* and *ERBB4* variability with reduced white matter density and integrity in human subjects ^21–23^.

In addition to the extensively studied *NRG1* and *ERBB4*, many other ligands and receptors in the ErbB signaling pathway are implicated in schizophrenia ^17^. However, the role of ErbB signaling in CNS myelination remains elusive due to the contradictory reports from different research groups ^24–27^. Heterozygous NRG1 type III mutant mice are reported to exhibit hypomyelination in the brain ^27^, whereas genetic ablation of all types of *NRG1* or *ErbB4* induces neither developmental alteration nor pathogenesis in the white matter of mutant mice ^24^. ERBB3 is the other ErbB receptor that specifically binds to the NRG family ligands and shows genetic association with schizophrenia ^28^. One research group reported that ErbB3 depletion in OLs from postnatal day 19 (P19) results in adult hypomyelination ^25^, whereas several other groups reported that ErbB3 knockout from embryonic age does not influence OLs and CNS myelin ^24, 26^.

Among four ErbB receptors, EGFR (i.e. ErbB1) binds specifically to the epidermal growth factor (EGF) family ligands ^19^. Notably, EGFR and its ligand EGF also exhibit genetic polymorphisms and aberrant expression that are associated with schizophrenia ^29–32^. Nevertheless, hypomorphic EGFR mutant mice display only delayed myelination in the brain, which is ascribed to impaired oligodendrogenesis during early development ^33^. Schizophrenia is a complex polygenic disorder. It is extremely noticeable that gene-gene interaction among variants in NRG-ERBB and EGF-ERBB pathways increases susceptibility to schizophrenia ^31, 34^. However, whether NRG-ErbB and EGF-ErbB signaling synergistically regulate myelin development and function has not been investigated.

In the CNS, OL precursor cells (OPCs) after terminal mitosis differentiate into newly-formed OLs (NFOs). NFOs progress from pre-myelinating OLs to newly myelinating OLs. Myelinating OLs effectively generate myelin sheaths in a short time window before further progressing into mature OLs (MOs) that maintain the myelin sheath ^35–40^. We demonstrated that, no matter which ErbB receptor is primarily bound by the ligand, both NRG-ErbB and EGF-ErbB receptors are active in OPCs and differentiated OLs. To study the oligodendroglial functions of ErbB signaling, we adopted an inducible pan-ErbB strategy that blocked the activities of any endogenous ErbB receptors in OL-lineage cells *in vivo*. By using two valuable research mouse tools that help distinguish the outcomes induced by defects of OLs at different differentiation stages, *Sox10*^+/rtTA^ (Tet-on) that target OPC-NFOs and *Plp*-tTA (Tet-off) that target MOs ^41^, we were able to demonstrate that disruption of ErbB signaling causes myelination-dependent and -independent white matter abnormalities, and prove that both pathophysiological changes lead to cognitive deficits.

## Materials and Methods

### Animals

*Plp*-tTA transgenic mice ^42^ were from the RIKEN BioResource Center (Stock No. RBRC05446). *Sox10*^+/rtTA^ mice were from Dr. Michael Wegner (University Erlangen-Nurnberg, Germany). Transgenic mice *TRE*-dnEGFR (Stock No. 010575) and *TRE*-ErbB2^V664E^ (Stock No. 010577) were from the Jackson Laboratory ^43^. Unless otherwise indicated, mice were housed under specific pathogen free conditions before experiments, in a room with a 12-hour light/dark cycle with access to food and water *ad libitum*. For biochemical and histological experiments, *Plp*-tTA;*TRE*-dnEGFR (*Plp*-dnEGFR), *Sox10^+/rtTA^; TRE*-dnEGFR (*Sox10*-dnEGFR), or *Sox10*^+/rtTA^;*TRE*-ErbB2^V664E^ (*Sox10*-ErbB2^V664E^) mice with either sex and their littermate control mice with matched sex were used. For indicated behavioral tests, only male mice were used, while both male and female mice were used for the rest behavioral tests because the results were not affected by sex difference. Tet-Off or Tet-On treatments were performed as previously reported ^41^.

### Electron Microscopy (EM)

Mice were anesthetized and transcardially perfused with 4% sucrose, 4% paraformaldehyde (PFA) and 2% glutaraldehyde in 0.1 mmol/L phosphate buffer (PB, pH 7.4). Different brain regions were isolated and post-fixed overnight at 4°C in 1% glutaraldehyde in 0.1 mmol/L PB. Samples were processed and ultrathin sections were obtained and stained with 2% uranyl acetate. EM images were captured with Tecnai 10 (FEI) and analyzed as previously reported ^41, 44^.

### Primary OPC culture and differentiation induction

The cortices of mice at P2–P4 were isolated and sheared into pieces, and digested by 0.25% Trypsin in Ca^2+^Mg^2+^-free D-Hanks’ balanced salt solution for 40 min at 37 °C. Trypsinized tissues were triturated and passed through 40 mm mesh. PDGFRα^+^ OPCs were obtained from the dissociated cells using MS Columns (Miltenyi Biotec 130-042-201) with magnetic beads (Miltenyi Biotec 130-101-547) according to the manufacturer’s recommendations. Purified OPCs were plated into poly-L-lysine-coated glass coverslips at a density of 25,000 cells per cm^2^ for immunofluorescence staining, or 35-mm dishes at 300,000 cells for western blotting, with DMEM/F12 plus 10% fetal bovine serum. Four hours later, the medium was replaced to DMEM/F12 with 2% B27, 20 ng/ml bFGF and 10 ng/ml PDGFAA. For differentiation induction of OPCs, the medium was replaced to DMEM/F12 with 2% B27 and 40 ng/ml triiodothyronine (T3). For analyzing the ErbB receptors that are activated by different ErbB ligands in OPCs or differentiated OLs, the medium was replaced to DMEM/F12 plus 2% B27 with HB-EGF (10 ng/ml, Peprotech, 100-47), EGF (20 ng/ml, Peprotech, 100-15) or NRG1 (100 ng/ml, Peprotech, 100-03), and cells were incubated at 37 °C with 5% CO_2_ for 10 min before cell lysis.

For OPC proliferation assay, the medium was replaced to DMEM/F12 plus 2% B27 with EGF (20 ng/ml) at day *in vitro* 1 (DIV1). At DIV2, cells were incubated with 10 μM EDU 6 h before fixation with 4% PFA, and processed for EDU staining with EdU Cell Proliferation Kit (Epizyme, CX004) and immunofluorescence staining. For OL differentiation assay, the medium was replaced to DMEM/F12 medium plus 2% B27 with EGF (20 ng/ml) or NRG1 (100 ng/ml) at DIV1, and cells were cultured for 4 days before fixation with 4% PFA for immunofluorescence staining. For cells from *Sox*-dnEGFR mice, 3 μg/ml Dox was added into the medium 24 h after plating.

### Immunofluorescence staining and TUNEL assay

Brain slices or coverslips were obtained and stained as previously described ^41, 45^. The primary antibodies used were: CC1 (1:500, Abcam, ab16794), NG2 (1:200, Abcam, ab50009), Ki67 (1:400, Cell Signaling Technology, 9129), TCF4 (1:500, Millipore, 05-511), Olig2 (1:500, Millipore, AB9610), MBP (1:100, Millipore, MAB382), ASPA (1:500, Oasis, OB-PRT005), PDGFRα (1:500, Oasis, OB-PRB011), Olig2 (1:3000, Oasis, OB-PGP040), MBP (1:2000, Abcam, ab7349). Apoptotic cells were examined with terminal deoxynucleotidyl transferase (TdT)-mediated deoxyuridine triphosphate (dUTP) nick-end labeling (TUNEL) assay according to the manufacturer’s instructions (Vazyme; Yeasen). Images were taken by a Zeiss LSM710 confocal microscope or a Nikon Eclipse 90i microscope. For cell counting based on immunostaining results, soma-shaped immunoreactive signals associated with a nucleus was counted as a cell.

### Western blotting

Subcortical white matter tissues were isolated and homogenized. Proteins were extracted and processed as previously reported ^41^. The primary antibodies used were: pErbB3 (1:2500, Abcam, ab133459), pErbB4 (1:2500, Abcam, ab109273), pErbB2 (1:2500, Abgent, AP3781q), EGFR (1:5000, Epitomics, 1902-1), pEGFR (1:2500, Epitomics, 1727-1), GAPDH (1:5000, Huabio, EM1101), MBP (1:1000, Millipore, MAB382), Olig2 (1:1000, Millipore, MABN50), Olig2 (1:1000, Huabio, ET1604-29), ErbB3 (1:200, Santa Cruz Biotechnology, sc-285), ErbB4 (1:200, Santa Cruz Biotechnology, sc-283), ErbB2 (1:200, Santa Cruz Biotechnology, sc-284), LDHA (1:2000, Abclonal, A1146), PKm2 (1:2000, Abclonal, A13905), GluT1 (1:2000, Abclonal, A6982), MCT2 (1:2000, Abclonal, A12386), MCT1 (1:2000, Abclonal, A3013), LDHB (1:2000, Abclonal, A18096). Intensities of protein bands were measured by Image J, and statistical analysis was performed after subtraction of the background intensity and normalization with controls in each batch of experiments to minimize the influences of batch-to-batch variations.

### *In situ* hybridization

Deeply anesthetized animals were perfused with 4% PFA in PBS, and isolated brains were post-fixed in 4% PFA in PBS at 4°C overnight. Following fixation, tissues were transferred to 20% sucrose in PBS overnight, embedded in OCT media, and then sectioned into 14 μm coronal sections on a cryostat sectioning machine (Thermo Fisher scientific, Microm HM525). RNA *in situ* hybridization was performed as previously described ^46^. To obtain mouse *Enpp6* riboprobes, a 1.3 kb fragment corresponding to Enpp6 mRNA (1400–2700 nt of NM_177304.4) was cloned into pBluescript II KS(-). The linearized plasmids were used as templates for *in vitro* transcription with T3 RNA polymerase (Promega) according to the manufacturer’s instructions.

### Real-time reverse transcription (RT)-PCR

Total RNA was extracted from isolated mouse white matter using TRIzol following manufacturer’s protocol. cDNA was synthesized by using the 5x All-In-One RT MasterMix (abmGood). Real-time PCR was performed in three repeats for each sample by using BrightGreen 2x qPCR MasterMix (abmGood) with the Bio-Rad CFX96 real-time PCR system as previously described ^43^. Relative mRNA levels were analyzed by software Bio-Rad CFX Manager 1.5. Transcripts of targeted genes were normalized to those of mouse 18S rRNA gene in the same samples. Primers for 18S rRNA were 5’-CGG ACA CGG ACA GGA TTG ACA and 5’-CCA GAC AAA TCG CTC CAC CAA CTA with a 94 bp PCR product. Primers for mouse EGFR gene and transgene dnEGFR were 5’-TCC TGC CAG AAT GTG AGC AG and 5’ -ACG AGC TCT CTC TCT TGA AG with a 500 bp PCR product. Primers for Enpp6 gene were 5’ - CAG AGA GAT TGT GAA CAG AGG C and 5’-CCG ATC ATC TGG TGG ACC T with a 123 bp PCR product. Primers for Slc12a2 gene were 5’-TTC CGC GTG AAC TTC GTG G and 5’ - TTG GTG TGG GTG TCA TAG TAG T with a 197 bp PCR product. Primers for Itpr2 gene were 5’-CCT CGC CTA CCA CAT CAC C and 5’-TCA CCA CTC TCA CTA TGT CGT with a 119 bp PCR product. Primers for GSN gene were 5’-ATG GCT CCG TAC CGC TCT T and 5’ - GCC TCA GAC ACC CGA CTT T with a 134 bp PCR product. Primers for Itgb4 gene were 5’ - GCA GAC GAA GTT CCG ACA G and 5’ - GGC CAC CTT CAG TTC ATG GA with a 157 bp PCR product. Primers for MBP gene were 5’-GGC GGT GAC AGA CTC CAA G and 5’-GAA GCT CGT CGG ACT CTG AG with a 167 bp PCR product. Primers for MAG gene were 5’ - CTG CCG CTG TTT TGG ATA ATG A and 5’-CAT CGG GGA AGT CGA AAC GG with a 127 bp PCR product. Primers for MOG gene were 5’-AGC TGC TTC CTC TCC CTT CTC and 5’-ACT AAA GCC CGG ATG GGA TAC with a 104 bp PCR product.

### RNA-Seq Analyses

Subcortical white matter tissues isolated from *Sox10*-dnEGFR and littermate controls, or *Sox10*-ErbB2^V664E^ mice and littermate controls, were used (three pairs for each group) for global transcriptome analysis by LC-Bio Co (Hangzhou, China). The final transcriptome was generated by Histat and StringTie. StringTie was used to estimate the expression level for mRNAs by calculating FPKM (Fragments Per Kilobase of exon model per Million mapped reads). Differentially expressed genes were identified by comparing FPKM of the mRNA reads from three sample pairs between *Sox10*-dnEGFR, or *Sox10*-ErbB2^V664E^ mice, and their littermate controls, by paired Student *t* test *via* MeV (MultiExperiment Viewer). Gene lists with significant difference (*P* < 0.05) in expression between *Sox10*-dnEGFR and littermate controls, or between *Sox10*-ErbB2^V664E^ and littermate controls, were compared, and genes with similar expression tendencies in *Sox10*-dnEGFR and *Sox10*-ErbB2^V664E^ mice were identified. Z value of these genes was calculated according to their FPKM by an equation “Z sample-i = [(log2(Signal sample-i)-Mean (Log2(Signal) of all samples)][Standard deviation (Log2(Signal) of all samples)]” and plotted as heat map by MeV. Gene Ontology (GO) term enrichment was analyzed by PANTHER Overrepresentation Test (Released 20171205) through http://geneontology.org with the significance estimated by Fisher’s Exact Test (FDR, false discovery rate).

### Behavioral Tests

Behavioral test paradigms were adopted from previous reports with modifications ^45, 47, 48^. Behavioral analyses for *Sox10*-dnEGFR and littermate controls with Dox-feeding from P21, or *Plp*-dnEGFR mice and littermate controls with Dox-withdrawal from P21, were carried out with 12- to 16-week-old animals by investigators unaware of their genotypes. Rotarod, pre-pulse inhibition (PPI), forced swim, and social interaction tests were performed as previously described ^48^. Tested mutant mice had littermate control mice with same sex. For PPI, social interaction, eight-arm radial water maze, Y maze, forced swim and tail suspension, all tested mice were male.

#### Open field and stereotyped behavior analysis

Animals were placed in a chamber (30 cm × 30 cm × 34.5 cm) and their movements were monitored and traced by a tracking software EthoVision XT 12 (Noldus, The Netherland). Locomotive activity was measured and summated at 5-min intervals over a 30-min period. Frequency and cumulative duration of stereotyped behaviors observed during 30-min traveling in the open field, including grooming, hopping, rearing supported and sniffing, were determined by EthoVision XT 12 and statistically analyzed. Anxiety of the animals was assessed by the differences of time that they spent in the central zone and peripheral zone during the 30 min.

#### Eight-arm radial water maze

According to the previous report with modification ^47^, animals were trained in eight-arm radial water maze for two weeks, with four trials each day to search a hidden platform in each trial for escaping from the water at 20-22 °C. Four hidden platforms were placed at the end of a same set of arms for all the training and tests, as illustrated in Supplementary Fig. 8C. After a trial that mice reached a hidden platform, mice were returned to their home cages with towel and warming pads. The visited platform was removed before the next trial. There was a 30-sec gap between trials. Mice aborted swimming during training were discarded. One day after training periods, the trained mice were tested for their working memory capacities that were represented by avoiding arms with visited platforms in previous trials. In the last trial of the test day, the animals had highest working memory load for they had to avoid swimming into three arms with platforms removed. First and repeat entries into any arm that previously had a platform were counted as working memory errors, and first entries into any arm that never had a platform were counted as reference memory errors which represent deficits in spatial recognition or long term memory. The day after test day, all the animals were tested in a simple visible platform task with 5 trials in a round pool, and each trial contained a visible platform placed at a different position 0.5-1.0 meter away from tested mice. The latency of mice reached the visible platform in each trial was averaged to assess their eyesight.

#### Y maze spontaneous alternation test

Animals were introduced in the center of a Y-shaped maze that have three arms (8 cm × 39 cm × 25 cm) at a 120° angle from each other, and their free movements in 8 min were monitored and traced by Anymaze software (Stoelting, Wood Dale, Illinois). The arm entries after 1 or 0 other arm visit were recorded as returned arm entries. The alternation errors were calculated by the percentage of returned arm entries in total arm entries. For the evaluation of the effect of L-lactate, same mice were tested one week later, with systemic L-lactate administration (i.p. 180 mg/kg) 2 h before the test.

#### Tail suspension

The tail suspension test was carried out 2 days later after the forced swim test, in which mice were suspended by using adhesive tape applied to the tail and videotaped for 5 min. Mouse movement during the 5 min was traced and analyzed by Anymaze software. For both forced swim and tail suspension tests, immobile period was defined by 70% of mouse bodies were motionless and lasted for at least 1 sec. Summation of immobile periods for each mouse was taken into statistical analysis.

### Electrophysiology

Following anesthesia and decapitation, optic nerves were isolated from mice and superfused with oxygenated artificial cerebrospinal fluid (ACSF) containing (in mmol/L): 119 NaCl, 2.5 KCl, 2.5 CaCl_2_, 1.3 MgCl_2_, 1.25 NaH_2_PO_4_, 26 NaHCO_3_, and 10 glucose; pH 7.4. Optic nerve CAP recording methods were adopted and modified from the previous reports ^49^. Briefly, two ends of the optic nerves were attached by suction electrodes, which backfilled with ACSF and connected to an IsoFlex (AMPI) for stimulation or a MultiClamp 700B (Molecular Device) for recording. The recorded signal was amplified 50 times, filtered at 30 kHz, and acquired at 20-30 kHz. Data were collected and analyzed by pClamp 10.3 software (Molecular Devices). The optic nerves were equilibrated for at least 30 min in the perfusion chamber in normal ACSF at room temperature before experiments. All experiments were performed at room temperature.

#### Maximal CAP recording

For each recorded nerve, stimulus pulse (100 μs duration) strength was adjusted with a stepped increase and finally to evoke the maximal CAP. CAPs were elicited 5 times at every step of the stimulus strength. After reaching the maximal CAP, the stimulus was increased an additional 25% for supramaximal stimulation to ensure the activation of all axons in the nerve. Note the supramaximal stimulation did not further change the CAPs. The areas under CAPs were calculated to determine the nerve conduction.

#### Oxygen-glucose deprivation (OGD) assay

The assay was performed as previously described with modification ^49, 50^. During experiments, CAPs were evoked by the supramaximal stimulus every 20 sec. After 60-min stimulation of the baseline CAP in normal condition, OGD was induced for the nerves by switching bathing solution from oxygenated ACSF (saturated with 95% O2/5% CO2) to glucose-free ACSF (replaced with equimolar sucrose to maintain osmolality) that was saturated with 95%N2/5%CO2. After 60-min OGD, oxygenated ACSF was restored and CAPs were recorded for another hour. CAPs recorded after 30-min baseline stimulation was taken as the initial CAPs. The effects of OGD on the nerve conduction and recovery were determined by normalizing the areas of CAPs recorded during OGD or recovery sessions to that of initial CAPs.

#### Neural activity dependence assay

The protocol was modified from published reports ^49, 50^. Before the experiments, CAPs were recorded every 30 sec to obtain baseline with the stimulus pulse strength set at the supramaximal levels. To evaluate the conduction of optic nerves under increasing energy demands, we gradually increased the stimulating frequency from 1 to 100 Hz. Each stimulating frequency was applied for 30-60 sec. For 1 and 5 Hz stimuli, CAPs were continuously recorded and CAP areas were measured for each CAP. For 10 to 100 Hz, nerves were stimulated by a train of 100 stimuli, and rest for 1 sec before the next train of stimuli. CAP areas were sampled for the last four stimuli of each train and averaged as one data point. For the statistical analysis, CAP areas were normalized to the initial levels. For L-lactate or D-lactate treatments, optic nerves were incubated with 20 mmol/L L-lactate or D-lactate in ACSF that contains 10 mmol/L glucose for 30 min before and during recording.

### Statistical analyses

Statistical analyses other than for RNA-seq data (described separately above) were performed using Prism (Graphpad) and presented as mean ± s.e.m.. Two-tailed unpaired Student’s *t* test was used for analysis between two groups with one variable, one-way ANOVA test was used for analysis among three or more groups with one variable, and two-way ANOVA test was used to determine difference among groups with two variables. Statistical significance was set at \**P* < 0.05, \*\**P* < 0.01, \*\*\**P* < 0.001.

## Results

### ErbB inhibition in oligodendrocytes at early differentiation stages induces hypomyelination

We characterized the expression of ErbB receptor members in subcortical white matter regions and found ErbB2 was barely detectable in mouse CNS myelin after P15 when most OPCs are differentiated, whereas EGFR, ErbB3 and ErbB4 were expressed with relatively stable levels during juvenile to adolescent development (Supplementary Fig. 1A, B). To analyze the receptors that are activated by NRG or EGF family ligands in OLs at different differentiation stages, we stimulated the purified OPCs and differentiated OLs, respectively, with NRG1 or EGF *in vitro*. Unexpectedly, tyrosine phosphorylation of ErbB1-4 receptors was all increased by either NRG1 or EGF stimulation (Fig. 1A, B). Even for ErbB2 that was expressed at very low levels in differentiated OLs, its phosphorylation upon EGF or NRG1 stimulation can still be detected (Fig. 1B). These results indicate that, irrespective of the stimulating ligands, all endogenous ErbB receptors in OPCs or OLs would exert synergistic effects. In other words, NRG-ErbB and EGF-ErbB signaling work together for the same biological processes. Therefore, the functions of ErbB signaling in OLs may not be accurately evaluated by investigating mice with knockout of only EGFR or ErbB3/4.

**Fig.1.**
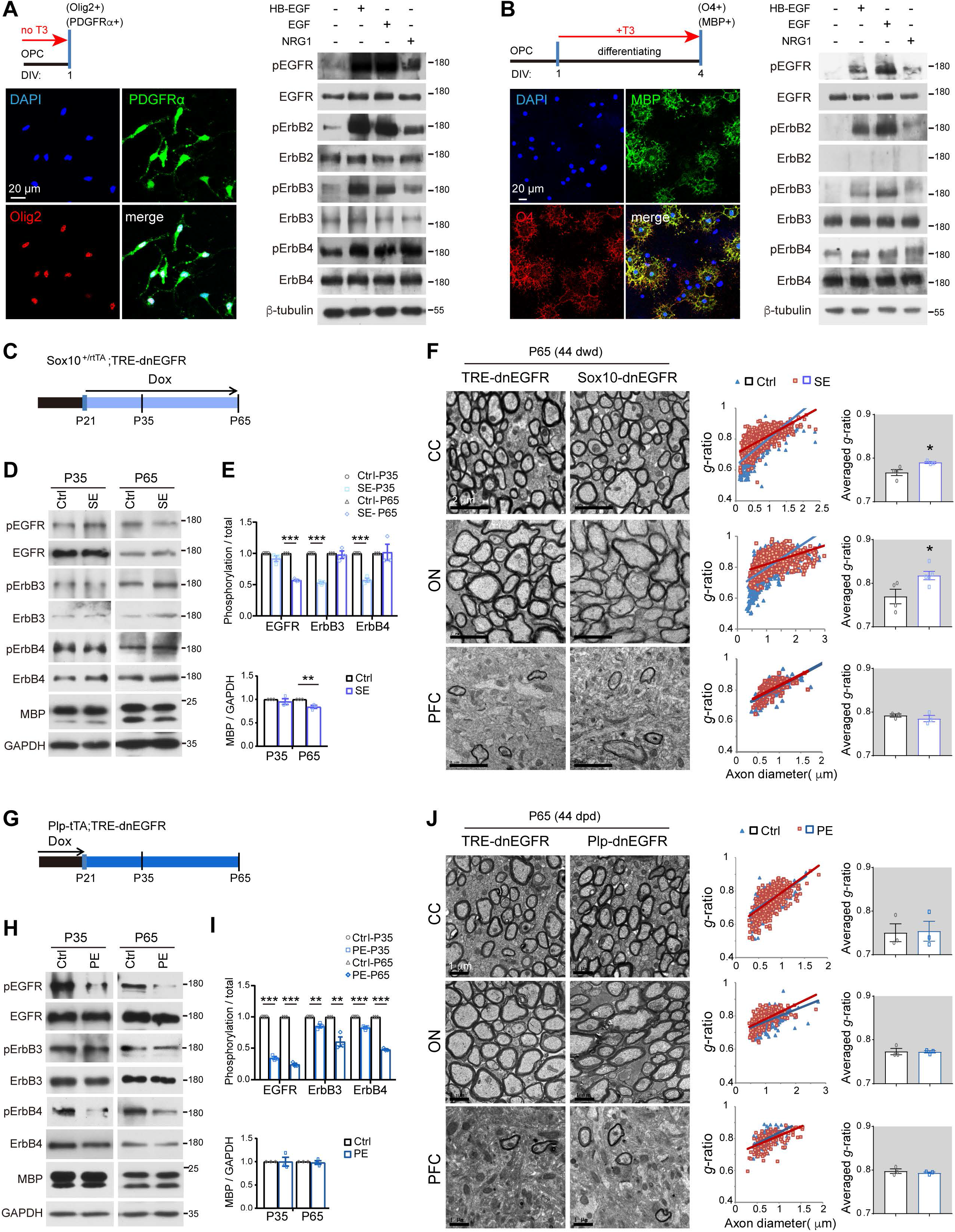
ErbB inhibition induces hypomyelination in *Sox10*-dnEGFR mice, but not in *Plp*-dnEGFR mice. **A**, **B** Activation of ErbB receptors by HB-EGF, EGF or NRG1 in OPCs (**A**) and differentiated OLs (**B**). The immunofluorescence results indicate the different OL stages. Activation status of each ErbB receptor was examined by western blotting with the specific antibody against its phosphorylated form. Note that ErbB2 receptor levels in differentiated OLs (**B**) were very low and hardly to be detected by the anti-ErbB2 antibody. **C**, **G** Dox treatment setting for indicated mice and littermate controls. **D**, **H** Indicated proteins and their activation status examined by western blotting of the white matter from *Sox10*-dnEGFR (SE) mice (**D**), or *Plp*-dnEGFR (PE) mice (**H**), in comparison with that of littermate controls (Ctrl). **E**, **I** Quantitative data of western blotting results. *n* = 3 for each group. **F**, **J** EM images of the corpus callosum (CC), optic nerve (ON), and prefrontal cortex (PFC) from *Sox10*-dnEGFR and littermate controls at 44 dwd (**F**), or *Plp*-dnEGFR and littermate controls at 44 dpd (**J**). *g*-ratio was calculated for myelinated axons and averaged *g*-ratio were analyzed by unpaired *t* test. In **F**, Ctrl *n* = 4, SE *n* = 3 for CC; Ctrl *n* = 4, SE *n* = 5 for ON; Ctrl *n* = 3, SE *n* = 3 for PFC. In **J**, *n* = 3 for each group.

To investigate the functions of synergistic ErbB signaling *in vivo*, we first generated *Sox10*^+/rtTA^;*TRE*-dnEGFR (*Sox10*-dnEGFR) mice (Supplementary Fig. 2A, B), an inducible mouse model that expresses a dominant negative mutant of EGFR (dnEGFR) in OPC-NFOs upon the treatment of doxycycline (Dox). DnEGFR is a truncated form of EGFR, losing the intracellular kinase domain but retaining the ability to form dimers with other ligand-bound ErbB members. When overexpressed, dnEGFR efficiently blocks the activation of any endogenous ErbB receptors under either NRG or EGF stimulation ^43^. In line with this concept, phosphorylation of ErbB3 and ErbB4 was reduced in white matter from *Sox10*-dnEGFR mice at P35 after 14 days with Dox feeding (dwd) (Fig. 1C-E). The phosphorylation of EGFR was not altered in *Sox10*-dnEGFR mice at P35 (Fig. 1D, E), consistent with our previous finding in *Sox10*-ErbB2^V664E^ mice that expressing a constitutively activated ErbB2 mutant in OPC-NFOs activates endogenous ErbB3 and ErbB4, but not EGFR ^41^.

Myelin thickness and ultrastructures in the white matter of *Sox10*-dnEGFR and littermate control mice at P35 with 14 dwd did not show significant differences (Supplementary Fig. 2C). Therefore, we raised these mice to adulthood with continuous Dox feeding. Phosphorylation of EGFR, instead of ErbB3 or ErbB4, was apparently reduced in the white matter of *Sox10*-dnEGFR mice at P65 (Fig. 1D, E). The change of dnEGFR targets in *Sox10*-dnEGFR mice from ErbB3/4 at P35 to EGFR at P65 may imply a switch of functional NRG-ErbB signaling to EGF-ErbB signaling in *Sox10*^+/rtTA^-targeted cells from adolescence to adulthood. For *Sox10*-dnEGFR mice at P65 with 44 dwd, axons in the corpus callosum and optic nerve were hypomyelinated (Fig. 1F). Consistently, myelin basic protein (MBP) was reduced in the white matter of *Sox10*-dnEGFR mice at P65 (Fig. 1D, E).

We further crossed *Plp*-tTA and *TRE*-dnEGFR to generate *Plp*-tTA; *TRE*-dnEGFR (*Plp*-dnEGFR) mice (Supplementary Fig. 2D, E). From embryonic age to P20, the expression of dnEGFR of *Plp*-dnEGFR was suppressed by Dox feeding. In these mice at P35 with dnEGFR expression in MOs for 14 days post Dox-withdrawal (dpd) at P21, western blotting revealed significant suppression on the phosphorylation of EGFR, as well as a mild suppression on that of ErbB3 and ErbB4 (Fig. 1G-I), consistent with their overactivation in *Plp*-ErbB2^V664E^ mice ^41^. No CNS myelin differences were observed in *Plp*-dnEGFR and littermate control mice at P35 with 14 dpd (Supplementary Fig. 2F).

We extended our investigation to P65 when dnEGFR still functionally suppressed ErbB receptor activities in the white matter of *Plp*-dnEGFR mice (Fig. 1H, I). The brains of *Plp*-dnEGFR and littermate mice at P65 with 44 dpd exhibited no difference in MBP protein levels (Fig. 1H, I) or myelin ultrastructures (Fig. 1J). These results suggest that the blockade of endogenous ErbB signaling in MOs does not affect CNS myelin development, whereas in OPC-NFOs induces hypomyelination.

### ErbB inhibition in oligodendrocytes at early differentiation stages increases oligodendrocyte numbers

To investigate whether hypomyelination is induced by myelinating cell number decreases, we examined the OL-lineage cells in *Sox10*-dnEGFR and *Plp*-dnEGFR mice. Unexpectedly, the densities of Olig2^+^ (all OL-lineage cells), NG2^+^ (OPCs), and CC1^+^ cells (post-mitotic OLs) significantly increased in the corpus callosum of *Sox10*-dnEGFR mice at P65 with 44 dwd (Fig. 2A, C), although the alteration was not detected at P35 with 14 dwd (Supplementary Fig. 3A, B). In contrast, no differences in Olig2^+^, CC1^+^, or NG2^+^ cell densities were observed in white matter of *Plp*-dnEGFR mice and littermate controls at either P35 or P65 (Fig. 2B, D and Supplementary Fig. 3C, D).

**Fig.2.**
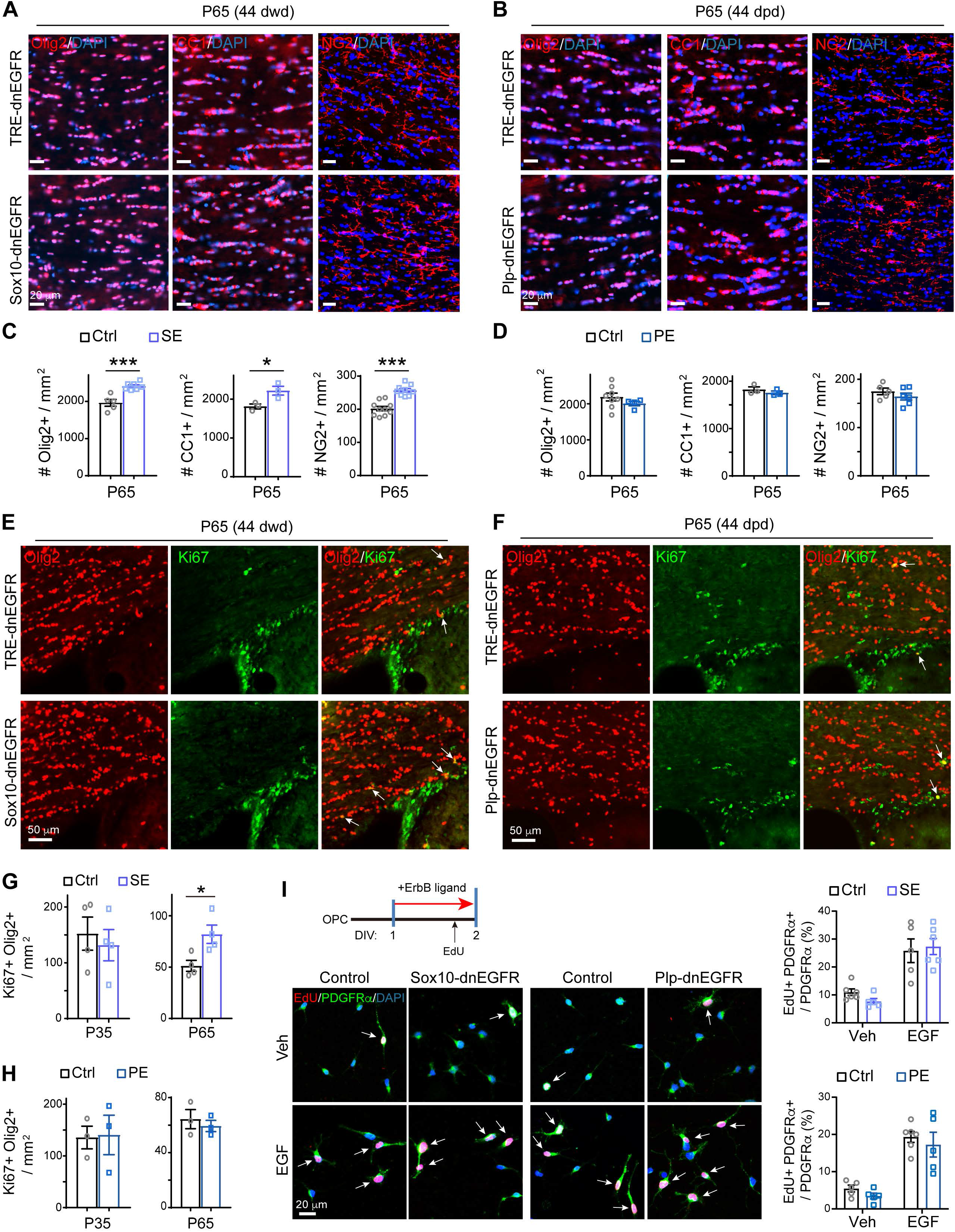
ErbB inhibition induces OL number increases in *Sox10*-dnEGFR mice, but not in *Plp*-dnEGFR mice. **A**, **B** Olig2^+^, CC1^+^, and NG2^+^ cells in the corpus callosum of indicated mice at indicated ages were examined by immunostaining. **C**, **D** Statistic results of Olig2^+^, CC1^+^, and NG2^+^ cell densities in the corpus callosum of *Sox10*-dnEGFR mice (SE) and littermate controls (Ctrl) at P65 with 44 dwd, or *Plp*-dnEGFR mice (PE) and littermate controls at P65 with 44 dpd. Data were from repeated immunostaining of 3 mice for each group. **E**-**H** Densities of proliferating OL lineage cells (Olig2^+^Ki67^+^) examined in indicated mice at indicated ages. Ctrl *n* = 4, SE *n* = 4 (**G**); Ctrl *n* = 3, PE *n* = 3 (**H**). **I** Proliferating OPCs (EDU^+^PDGFRα^+^) examined in OPCs purified from indicated mice. For SE OPCs in comparison with control OPCs (top graph): Ctrl with vehicle treatment (veh) *n* = 6, SE with veh *n* = 5; Ctrl with EGF treatment (EGF) *n* = 5, SE with EGF *n* = 6. For PE OPCs in comparison with control OPCs (bottom graph): Ctrl with veh *n* = 5, PE with veh *n* = 5; Ctrl with EGF *n* = 6, PE with EGF *n* = 5.

OL number increases were due to the increased proliferating OPCs (Ki67^+^Olig2^+^) that were detected in the white matter of *Sox10*-dnEGFR mice at P65 (Fig. 2E, G and Supplementary Fig. 4A), where apoptotic cells (TUNEL^+^) were as minimal as that in control mice (Supplementary Fig. 4C). In contrast, *Plp*-dnEGFR white matter that does not have OL number changes exhibited no changes in Ki67^+^Olig2^+^ or TUNEL^+^ status (Fig. 2F, H and Supplementary Fig. 4B, D). However, ErbB signaling does not negatively regulate OPC proliferation, as we found that EGF increased the numbers of EdU^+^ OPCs *in vitro* (Fig. 2I). We have reported that *Sox10*^+/rtTA^ mostly targets terminally differentiating OPCs ^41^. Consistent with that ErbB signaling in proliferating OPCs was not disrupted, neither *Plp*-dnEGFR nor *Sox10*-dnEGFR OPCs had lost the proliferating responses to EGF (Fig. 2I). Therefore, enhanced OPC proliferation and increased OL numbers in adult *Sox10*-dnEGFR mice does not seemed to be directly induced by ErbB disruption in OPCs, but rather an indirect consequence through cell communications during brain development.

### ErbB inhibition suppresses NFO differentiation to lead to hypomyelination

With the OL number increased, there must be other events linking to myelination capacity are impaired in the brain of *Sox10*-dnEGFR mice. Given the fact that *Sox10*-dnEGFR and *Sox10*-ErbB2^V664E^ mice both exhibited hypomyelination ^41^, they should share a molecular or cellular deficit in myelination. We performed RNA-seq analyses of subcortical white matter tissues and identified 68 genes which exhibited similar expression tendencies in *Sox10*-ErbB2^V664E^ and *Sox10*-dnEGFR mice during adolescence (Fig. 3A, Supplementary Fig. 5A-D, and Supplementary Tables 1 and 2). This group of genes have potential to regulate CNS myelin development. Notably, in addition to *Gsn* and *Itgb4* that are characteristic genes for myelinating OLs ^51^, *Enpp6, Itpr2*, and *Slc12a2* as characteristic genes for NFOs also exhibited significantly reduced expression in both mouse lines. Transcription of these genes were persistently reduced in *Sox10*-dnEGFR white matter at P65 (Fig. 3B). Although the NFO numbers became very few and the differences became indistinguishable for mice at P65, by *Enpp6 in situ* hybridization ^37^ and TCF4 immunostaining ^52–54^ that specifically label NFOs, we confirmed the deficiency of NFOs in *Sox10*-dnEGFR mice at P35 (Fig. 3C, D and Supplementary Fig. 6A). In support of the notion that ErbB signaling is required for NFO differentiation, in an *in vitro* assay that covers NFO differentiation process, *Sox10*-dnEGFR NFOs lost the responses to EGF and NRG1 in generating MBP^+^ myelinating OLs (Fig. 3E, F). The progression of myelinating NFOs into MOs is not altered by *Sox10*-dnEGFR, because with MO (ASPA^+^) numbers increased in *Sox10*-dnEGFR mice at P65, the ratio of MOs in total OLs (ASPA^+^/Olig2^+^) was the same in *Sox10*-dnEGFR as in littermate control mice (Fig. 3G, H). In spite the increased MO numbers, the transcription of myelin proteins, *Mbp, Mag*, and *Mog*, were reduced in *Sox10*-dnEGFR at P65, but not in P35 (Fig. 3B), consistent with the observation by EM. Therefore, hypomyelination in *Sox10*-dnEGFR mice is attributed to the NFO deficiency, where NFO differentiation was impaired shortly after ErbB inhibition, although it took 44 days to result in obvious hypomyelination (Fig. 3I). In contrast, the *in vivo* transcription of NFO characteristic genes and myelin protein genes, the ASPA^+^ MO numbers, as well as the *in vitro* MBP^+^ myelinating NFOs induced by ErbB ligands, were not altered in *Plp*-dnEGFR mice (Fig. 3B, E, F and Supplementary Fig. 6B-D).

**Figure 3.**
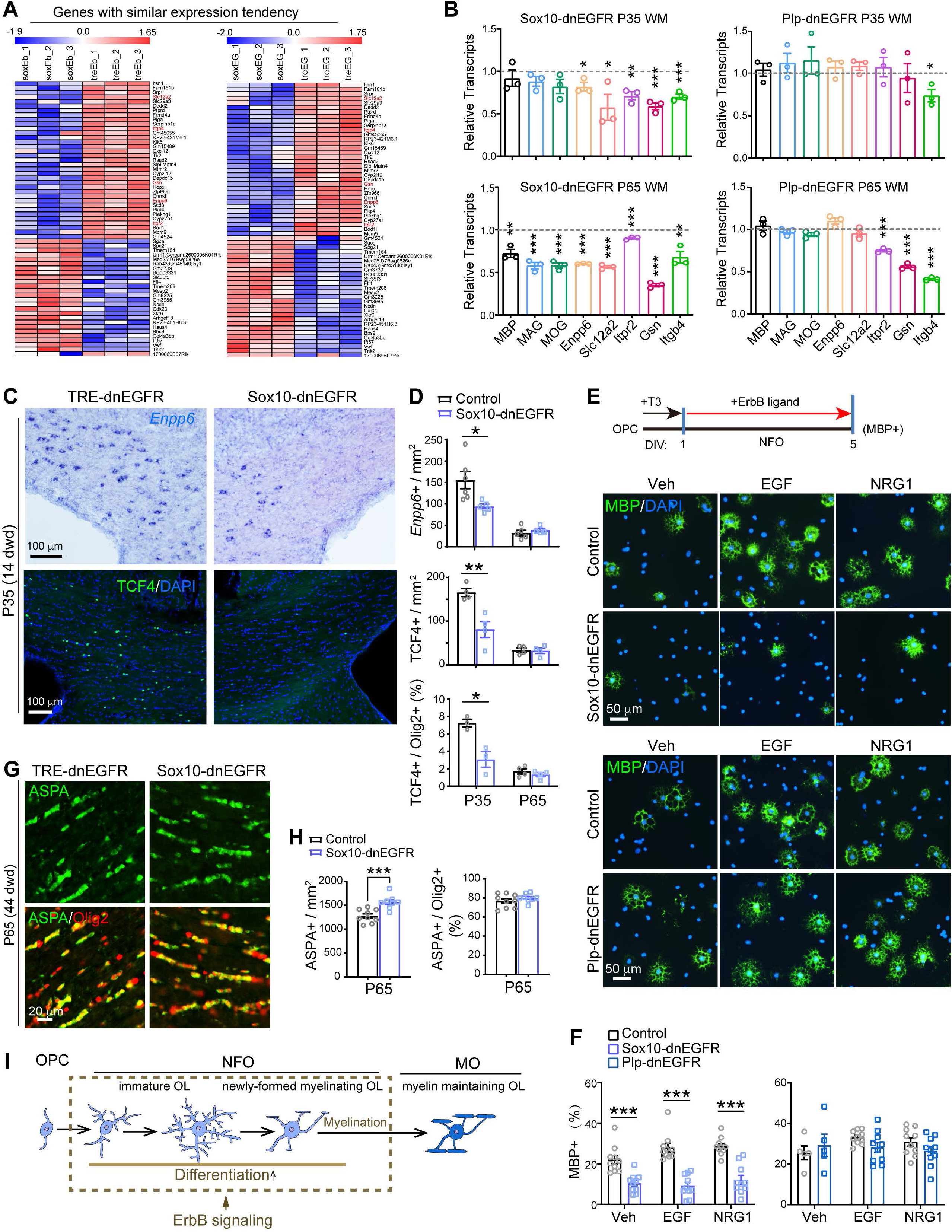
ErbB inhibition impairs NFO differentiation during adolescent development. **A** RNA-seq was performed for white matter tissues isolated from *Sox10*-ErbB2^V664E^ mice at P30 with 9 dwd, or *Sox10*-dnEGFR mice at P35 with 14 dwd, and their littermate controls. For each mouse group we sequenced three pairs of samples and identified 2298 genes that had altered expression in the white matter of *Sox10*-ErbB2^V664E^ mice (Supplementary Table 1), as well as 1184 genes in that of *Sox10*-dnEGFR mice (Supplementary Table 2). By comparing the two groups of genes, 68 genes with similar expression tendencies in the white matter of *Sox10*-ErbB2^V664E^ (soxEb *vs* treEb) and *Sox10*-dnEGFR (soxEG *vs* treEG) mice were identified. Heat maps of Z value of each gene are presented. The detailed RNA-seq data have been deposited in the GEO and SRA database and can be found at GEO: GSE123491. **B** Transcription levels of characteristic genes of OLs at different differentiation stages in the white matter (WM) of indicated mice. Shown are real-time RT-PCR results of indicated mice that have been normalized by those of littermate controls. Mouse pairs for each group *n* = 3. **C** *In situ* hybridization results of *Enpp6* and immunostaining results of TCF4 in the corpus callosum of *Sox10*-dnEGFR mice and littermate controls at indicated ages. **D** Statistic analyses of *Enpp6^+^* or TCF^+^ cell densities, as well as the ratios of NFOs in total OLs (TCF4^+^/Olig2^+^) in indicated mice at indicated ages. Data were from repeated immunostaining of 3 mice for each group. **E, F** Newly-myelinating OLs (MBP^+^) induced by EGF or NRG1 after differentiation induction of OPCs from indicated mice with triiodothyronine (T3). For analysis of *Sox10*-dnEGFR NFO differentiation (left in **F**): Control with vehicle treatment (veh) *n* = 10, *Sox10*-dnEGFR with veh *n* = 10; Control with EGF treatment (EGF) *n* = 10, *Sox10*-dnEGFR with EGF *n* = 10; Control with NRG1 treatment (NRG1) *n* = 10, *Sox10*-dnEGFR with NRG1 *n* = 10. For analysis of *Plp*-dnEGFR NFO differentiation (right in **F**): Control with veh *n* = 5, *Plp*-dnEGFR with veh *n* = 5; Control with EGF *n* = 10, *Plp*-dnEGFR with EGF *n* = 10; Control with NRG1 *n* = 10, *Plp*-dnEGFR with NRG1 *n* = 10. **G** Immunostaining results of MOs (ASPA^+^Olig2^+^) in the corpus callosum of *Sox10*-dnEGFR and littermate controls at indicated ages. **H** Statistic results of ASPA^+^ cell densities and the ratio of ASPA^+^ to Olig2^+^ cell numbers in indicated mice at indicated ages. Data were from repeated immunostaining of 3 mice for each group. **I** Schematic illustration of OL differentiation stages, among which oligodendroglial ErbB signaling regulates NFO differentiation.

### ErbB inhibition in MOs disrupts cognitive function in the absence of myelin alteration

It is noticeable that, although *Plp*-dnEGFR white matter did not exhibit alteration in the expression of myelin protein genes and NFO genes, it exhibited reduction in the transcripiton of *Itpr2, Gsn*, and *Itgb4* (Fig. 3B). These results suggested that OLs in *Plp*-dnEGFR mice were not as normal as we assumed. Before the mechanistic investigation, we first evaluated whether ErbB signaling in MOs is important for any brain functions by comparing the behavioral performance of *Sox10*-dnEGFR and *Plp*-dnEGFR mice.

With minimal differences in the PNS (Supplementary Fig. 7A-F), *Sox10*-dnEGFR mice performed worse than control mice in the rotarod test (Fig. 4A), and were slightly hypoactive in the open field test (Fig. 4B). Nevertheless, they performed normally in the central/peripheral zone analysis for assessment of anxiety, stereotyped behavior analysis and social interaction analysis for potential autistic-like phenotype, pre-pulse inhibition analysis for sensory gating, as well as forced swim and tail suspension tests for depression (Fig. 4E, F, I, M and Supplementary Fig. 8A). Interestingly, *Plp*-dnEGFR mice performed normally, similar to the controls in most tests, except exhibiting a subtle hyperactivity in the open field test (Fig. 4C, D, G, H, J, N and Supplementary Fig. 8B). The different results from the two mouse models implied that the impaired motor coordination is attributed to hypomyelination in the CNS.

**Fig.4.**
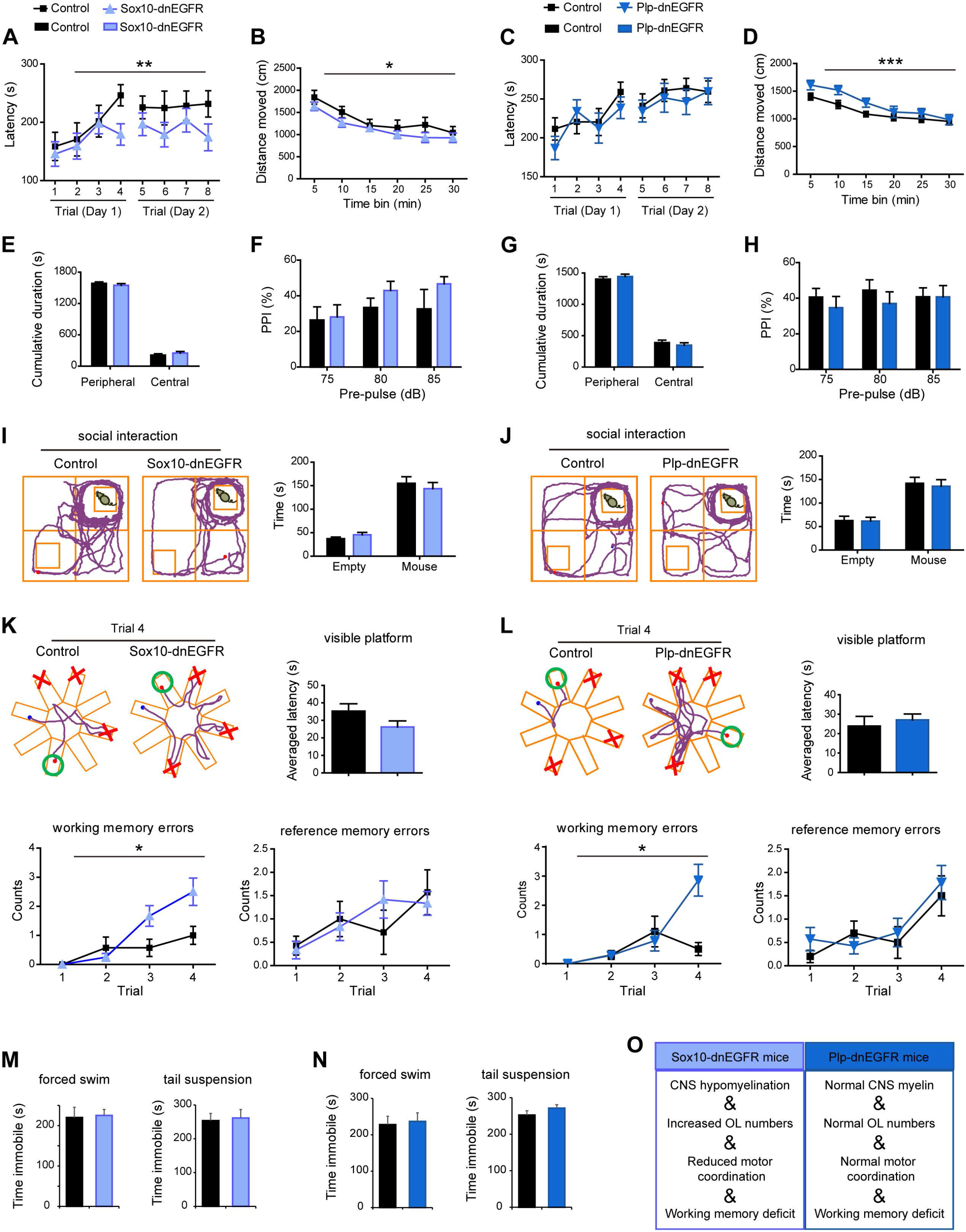
ErbB inhibition in MOs impairs working memory in the absence of myelin alteration. Behavioral performance of adult *Sox10*-dnEGFR mice with littermate controls, or *Plp*-dnEGFR mice with littermate controls. **A**, **C** Motor coordination assessed by rotarod test: controls *n* = 12, *Sox10*-dnEGFR *n* = 12 (**A**); control *n* = 13, *Plp*-dnEGFR *n* = 12 (**C**). **B**, **D** Locomotive activity assessed by open field tests: control *n* = 11, *Sox10*-dnEGFR *n* = 13 (**B**); control *n* = 19, *Plp*-dnEGFR *n* = 14 (**D**). **E**, **G** Zone analysis of open field tests: control *n* = 11, *Sox10*-dnEGFR *n* = 13 (**E**); control *n* = 19, *Plp*-dnEGFR *n* = 14 (**G**). **F**, **H** Sensory gating assessed by PPI tests: control *n* = 10, *Sox10*-dnEGFR *n* = 12 (**F**); control *n* = 14, *Plp*-dnEGFR *n* = 12 (**H**). **I**, **J** Social interests assessed by social interaction tests: control *n* = 12, *Sox10*-dnEGFR *n* = 13 (**I**); control *n* = 14, *Plp*-dnEGFR *n* = 13 (**J**). **K, L** Working memory assessed by eight-arm radial water maze test: control *n* = 7, *Sox10*-dnEGFR *n* = 12 (**K**); control *n* = 10, *Plp*-dnEGFR *n* = 14 (**L**). **M**, **N** Desperation assessed by forced swim and tail suspension tests: control *n* = 10, *Sox10*-dnEGFR *n* = 13 (**M**); control *n* = 13, *Plp*-dnEGFR *n* = 13 (**N**). Data were analyzed by two-way ANOVA test, except for those from the forced swim test, tail suspension test, and visible platform test in eight-arm radial water maze that were analyzed by unpaired *t* test. Illustrative examples of the travel pathways of control or mutant mice in social interaction and eight-arm radial water maze were included in **I**, **J**, **K** and **L**. In illustrative examples of eight-arm radial water maze in **K** and **L**, green circles indicate the last arms with a hidden platform, while red crosses indicate the arms with used platforms in the past three trials. **O** Summary of the phenotypes of *Sox10*-dnEGFR and *Plp*-dnEGFR mice.

We further tested these mice in the eight-arm radial water maze, a paradigm analyzing working memory capacity. Not only *Sox10*-dnEGFR mice, which had CNS hypomyelination, but also *Plp*-dnEGFR mice, which did not have myelin alteration, had significantly more working memory errors than control mice (Fig. 4K, L and Supplementary Video 1). Note that they had normal eyesight as performed in the visible platform test, as well as similar reference memory errors that indicated unaltered spatial recognition and memory (Fig. 4K, L). This phenotype in *Plp*-dnEGFR mice reveals that working memory deficiency can be caused directly by ErbB inhibition in MOs through a myelination-independent mechanism.

### ErbB inhibition in MOs suppresses axonal conduction under energy stress

To determine what kind of function was impaired in white matter tracts of *Plp*-dnEGFR mice, we acutely isolated the optic nerves from adult mice and recorded electrical stimulus-evoked compound action potentials (CAPs). The maximal CAPs, which represent that all axons in the nerves are excited, were similar in *Plp*-dnEGFR optic nerves and control nerves (Fig. 5A). In contrast, they were reduced in *Sox10*-dnEGFR optic nerves (Fig. 5B). These results reflected that the basic axonal conduction was not affected in *Plp*-dnEGFR white matter tracts, whereas it was impaired in *Sox10*-dnEGFR white matter tracts that exhibited hypomyelination.

**Fig.5.**
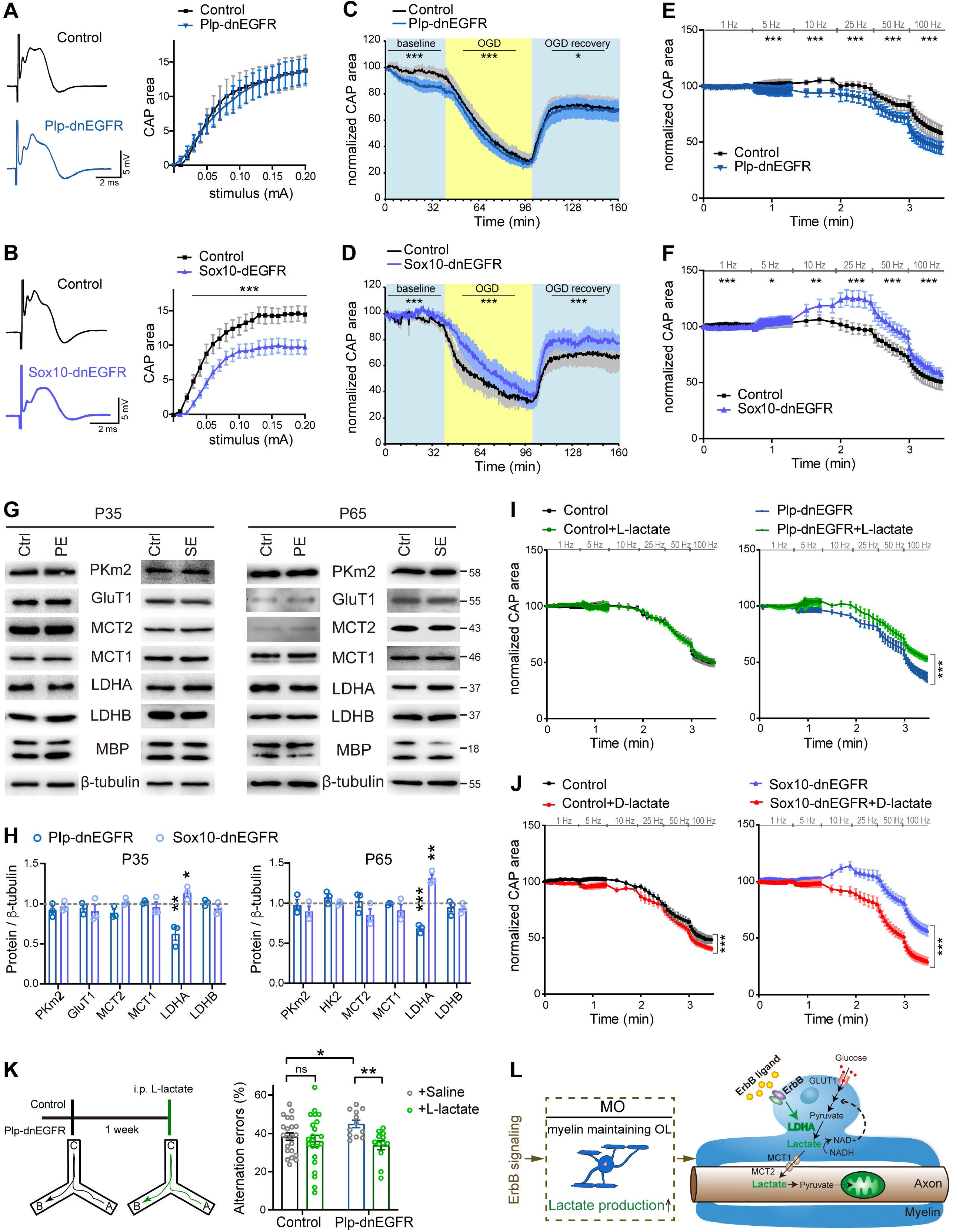
ErbB inhibition in MOs impairs lactate production, suppressing axonal conduction under energy stress. **A**, **B** Axonal excitability is similar in *Plp*-dnEGFR optic nerves and control nerves (**A**), but decreased in *Sox10*-dnEGFR optic nerves in comparison with controls (**B**). CAPs of optic nerves generated by electrical stimuli with intensities at stepped increase (0-0.2 mA) were recorded *ex vivo*. Data were from 3-7 optic nerves of 3-5 mice for each group. **C**, **D** The CAP decline induced by OGD is slightly accelerated and aggravated in *Plp*-dnEGFR optic nerves (**C**), but decelerated and attenuated in *Sox10*-dnEGFR optic nerves (**D**). OGD was started for the recorded nerves after 1-h baseline stimulation, and stopped after another hour by restoring the bathing media to oxygenated ACSF. Initial CAPs were recorded after 30-min baseline stimulation. The areas under CAPs were measured and normalized to the initial levels. Data were from 4-8 optic nerves of 3-5 mice for each group. **E**, **F** Neuronal activities with frequency at 5-100Hz accelerated the CAP decline in *Plp*-dnEGFR optic nerves (**E**), but increased the CAPs at 10-25Hz and slowed the CAP decline at 50-100Hz in *Sox10*-dnEGFR optic nerves (**F**), in comparison with their controls, respectively. Data were from 4-8 optic nerves of 3-5 mice for each group. **G** Indicated proteins examined by western blotting in the white matter of *Sox10*-dnEGFR (SE) mice or *Plp*-dnEGFR (PE) mice, in comparison with that of littermate controls (Ctrl). **H** Quantitative data of western blotting results. *n* = 3 for each group. **I** L-lactate restored the declined CAPs in *Plp*-dnEGFR optic nerves. Data were from 6-10 optic nerves of 3-5 mice for each group. **J** D-lactate suppressed the expanded CAPs in *Sox10*-dnEGFR optic nerves. Data were from 5-12 optic nerves of 3-6 mice for each group. **K** Working memory deficits in *Plp*-dnEGFR mice were rescued by systemic L-lactate as assessed by the Y maze test. Control with saline or L-lactate *n* = 22; *Plp*-dnEGFR with saline or L-lactate *n* = 12. **L** Schematic illustration of the mechanism that ErbB signaling promotes aerobic glycolysis in MOs.

In addition to myelin formation, OLs also offer essential trophic support to neurons, especially supporting axonal energy consumption under energy stress ^55^. Next, we challenged the optic nerves by incubating them in the oxygen-glucose deprivation (OGD) condition for 60 min. CAPs fell gradually in control optic nerves, and finally fell to 30% of the initial levels (Fig. 5C, D). However, for *Plp*-dnEGFR optic nerves in the OGD condition, CAP failure was slightly accelerated and aggravated (Fig. 5C). In contrast, for *Sox10*-dnEGFR optic nerves under the same condition, CAP failure was decelerated and attenuated (Fig. 5D). When the glucose and oxygen levels in the bathing medium were restored, CAPs in control optic nerves and *Plp*-dnEGFR optic nerves recovered to 60% of the initial levels (Fig. 5C, D). However, in *Sox10*-dnEGFR optic nerves, CAPs recovered to 80% of the initial levels (Fig. 5D).

It is notable that continuous electrical stimulation caused a baseline CAP decline in *Plp*-dnEGFR optic nerves, whereas a baseline CAP enhancement in *Sox10*-dnEGFR optic nerves, before the OGD (Fig. 5C, D). Therefore, we further examined the axonal conduction under a physiological condition with increasing energy demands generated by neuronal activities ^49, 50^. By stimulating the optic nerves with several trains of short bursts with frequency increased from 1 to 100 Hz, the 5-25Hz electrical stimuli amplified CAPs and the 50-100Hz stimuli induced smaller CAP decay in *Sox10*-dnEGFR optic nerves (Fig. 5F). In contrast, in *Plp*-dnEGFR optic nerves, either group of stimuli significantly aggravated the CAP decay (Fig. 5E).

### ErbB inhibition in MOs suppresses its lactate supply to electrically active axons

Note that MO numbers were not altered in *Plp*-dnEGFR mice that have ErbB inhibition in MOs, whereas ErbB receptors were not inhibited in MOs of *Sox10*-dnEGFR mice that have increased MOs (Fig. 4O). Therefore, the opposite CAP maintaining capacities of *Sox10*-dnEGFR and *Plp*-dnEGFR optic nerves in the energy-challenging studies reveal that ErbB signaling in MOs determines the glia-axon energy coupling efficiency. In support of this notion, we discovered a significant decrease of lactate dehydrogenase A (LDHA), the LDH subunit that promotes lactate production, in *Plp*-dnEGFR white matter (Fig. 5G, H). The other enzymes and transporters in glycolysis pathways coupled between OLs and axons, including PKm2, GluT1, LDHB, MCT1, and MCT2, were not altered (Fig. 5G, H). Interestingly, *Sox10*-dnEGFR white matter exhibited increased LDHA (Fig. 5G, H), in line with its resistance to energy stress. These results support the idea that lactate is the main energy substrate that MOs provide to axons. Consistent with this idea, supplementation of L-lactate rescued the accelerated CAP decline in *Plp*-dnEGFR optic nerves upon high-frequency stimulation (Fig. 5I). On the other hand, the expanded CAPs in *Sox10*-dnEGFR optic nerves was suppressed by D-lactate (Fig. 5J), the metabolically inert isoform that competes for the endogenous lactate uptake and inhibits LDH activity ^50, 56^. Futhermore, L-lactate supplementation through systemic circulation rescued working memory deficits in *Plp*-dnEGFR mice (Fig. 5K). Of note, the administration of L-lactate in control mice exhibited no statistic significance on improving axonal conduction or cognition (Fig. 5I, K). The results indicate that endogenous lactate produced by aerobic glycolysis in normal white matter tracts is saturated, further emphasizing the importance of oligodendroglial ErbB signaling in lactate production.

## Discussion

There have been genetic variants revealed in schizophrenic patients that are associated with reduced expression or loss of function in ErbB receptors or ligands ^19, 20, 57, 58^. The present study reveals that the loss of ErbB functions in OLs have two types of adverse impacts on white matter tracts: hypomyelination and deficient lactate generation. By using the mouse models *Sox10*-dnEGFR and *Plp*-dnEGFR, which help distinguish the consequences induced by ErbB disruption in OPC-NFOs from that in MOs, we demonstrated that ErbB inhibition in OPC-NFOs impairs NFO differentiation, resulting in hypomyelination in adulthood. Simultaneously, ErbB inhibition in MOs blocks the supply of lactate to axons, disrupting axonal conduction within electrically active neural circuits. Importantly, both of the defects contributed to working memory deficits, firmly supporting that both myelination-dependent and -independent oligodendropathy can be primary causes to the cognitive symptoms of schizophrenia.

Our results demonstrated that both ErbB3/ErbB4 receptors binding to the NRG family ligands and EGFR binding to the EGF family ligands are functional in adolescent and adult OLs, and ErbB3/4 are more active than EGFR in adolescent OPC-NFOs (Fig. 1D, E). In spite of the contradictory findings from different research groups on the studies of ErbB3 and ErbB4 mutant mice, the phenotypes observed in *Sox10*-dnEGFR mice are consistent with the previous report in which ErbB3 depletion from P19 results in adult hypomyelination and working memory deficits ^25^. Beyond this, the present study elaborates the pro-myelinating role of ErbB signaling in NFOs (Fig. 3A-I).

The phenotypes exhibited in *Sox10*-dnEGFR mice provide the direct evidence supporting an known concept that myelin integrity is fundamental to cognitive performance of patients ^59^. In contrast, *Plp*-dnEGFR mice are a good model to affirm the myelination-independent contributions of OLs to higher brain function, given that dual inhibition of NRG-ErbB and EGF-ErbB signaling in MOs does not affect myelin or OL numbers in the adolescent and adult brains (Figs. 1 and 2). Our findings from *Plp*-dnEGFR mice emphasize the importance of lactate production in myelin to cognition (Figs. 4 and 5). Strategies improving lactate in the brain may combat the deficits induced by glia-axon energy coupling deficiency, as suggested by our findings that supplementation of L-lactate restored the axonal conduction and working memory capacity in *Plp*-dnEGFR mice (Fig. 5I,K). Lactate generated by exercise has been shown to improve learning and memory ^60^. Therefore, the present study may suggest a biological basis for the beneficial roles of sports for psychiatric patients to ameliorate cognitive deficits ^61, 62^.

ErbB signaling has been revealed to regulate aerobic glycolysis in cancer cells, where the expression of PKm2, LDHA, and GluT1, are increased to optimize cancer cells for the Warburg effect ^63–65^. It was unexpected that ErbB signaling in MOs are indispensable for LDHA expression only (Fig. 5G, H), highlighting a cell specific event regulated by ErbB signaling. Importantly, convertion of pyruvate to lactate regenerates NAD+, which fuel the conversion of glyceraldehyde-3-phosphate to 1,3-biophosphoglycerate and secure the constantness of aerobic glycolysis ^66^. Therefore, ErbB-regulated lactate generation in MOs maintains the axonal conduction under energy stress, meanwhile accelerates the flux of glycolysis that provides multiple intermediates for biosyntheses (Fig. 5L). It is worthy of further investigation to evaluate the full functions of ErbB signaling in MOs.

Given the fact that axonal conduction under energy stress was enhanced in *Sox10*-dnEGFR white matter tracts (Fig. 5D, F), but *Sox10*-dnEGFR mice still exhibited working memory deficits (Fig. 4K), structural integrity of myelin outweighs the lactate production capacity in determination of cognitive performance. Targeting NFO differentiation may be more important in treatments to combat white matter abnormalities induced by ErbB dysregulation. However, severe and long-term deficits of OL metabolite supply also cause axonal damage and myelin degeneration ^67, 68^. Moreover, myelin is very sensitive to environmental insults. Modest disruption of ErbB signaling by genetic mutation may be able to render myelin vulnerable to such insults, aggravating focal loss of connections under conditions of stress, ischemia, sleeplessness, trauma, etc. Note that a recent study has associated glucose metabolic disturbance with white matter structural abnormalities in drug-naive schizophrenia ^69^. Therefore, OL metabolic dysfunction in patients may eventually evolve into a detectable structural alteration in white matter that contributes to brain dysfunctions other than working memory deficits. Whether modest ErbB disruption increases the vulnerability of white matter to environmental etiological factors warrants further investigation.

## ACKNOWLEDGMENTS

We thank Zhengdong Wei, Xiaoyan Lu, Kaiwei Zhang, Youguang Yang, and Shasha Zhang in Hangzhou Normal University for the assistance in EM image analyses, and Wanhua Shen for the assistance in electrophysiological experiments. This work was supported by the ministry of science and technology of China (2021ZD0201700 to Y.T.), the national natural science foundation of China (31371075, 31871030 and 32170956 to Y.T.), the key project of Zhejiang provincial natural science foundation of China (2022XHSHH004 to J.J.), and the young scientists project of Zhejiang provincial natural science foundation of China (LQ19H090009 to J.J.).

## AUTHOR CONTRIBUTION

Y.T. conceived, designed and supervised research; X.H., Q.Z., T.L., Q.H., H.L., X.N., L.H., H.H, and Y.X. performed experiments; Q.M., Y.S., and J.J. provided technique supports; X.H., Q.Z., T.L., Q.H., H.L., and Y.T. analyzed data; Y.T. wrote the paper.

## ADDITIONAL INFORMATION

Supplementary Information, including Supplementary Figures 1-8, Supplementary Tables 1-2, and Supplementary Video 1, is available at MS’s website.

