## Supplementary for "ErbB inhibition impairs cognition via disrupting myelination and aerobic glycolysis in oligodendrocytes"

**Running title: ErbB regulates myelination and glial glycolysis**

**Authors**

Xu Hu,[1,2,4] Qingyu Zhu,[2] Tianjie Lou,[2] Qianqian Hu,[2] Guanxiu Xiao,[2] Huashun Li,[2] Xiaojie Niu,[2] Li He,[2] Hao Huang,[2] Yifei Luan,[2] Yijia Xu,[2] Mengsheng Qiu,[2] Ying Shen,[3] Jiemin Jia,[4] and Yanmei Tao,[2,5,*]

**Affiliations**

[1]College of Life Sciences, Zhejiang University, Hangzhou 310058, China

[2]Key Lab of Organ Development and Regeneration of Zhejiang Province, Institute of Life Sciences, College of Life and Environmental Sciences, Hangzhou Normal University, Hangzhou 311121, China.

[3]Department of Neurobiology, Key Laboratory of Medical Neurobiology of Zhejiang Province, Zhejiang University School of Medicine, Hangzhou 310058, China.

[4]Key Laboratory of Growth Regulation and Translational Research of Zhejiang Province, School of Life Sciences, Westlake University, Hangzhou 310024, China

[5]Department of Physiology, Medical College, Southeast University, Nanjing 210009, China.

**Supplementary Information** includes Supplementary Figures 1-8, Supplementary Tables 1-2, and Supplementary Video 1.

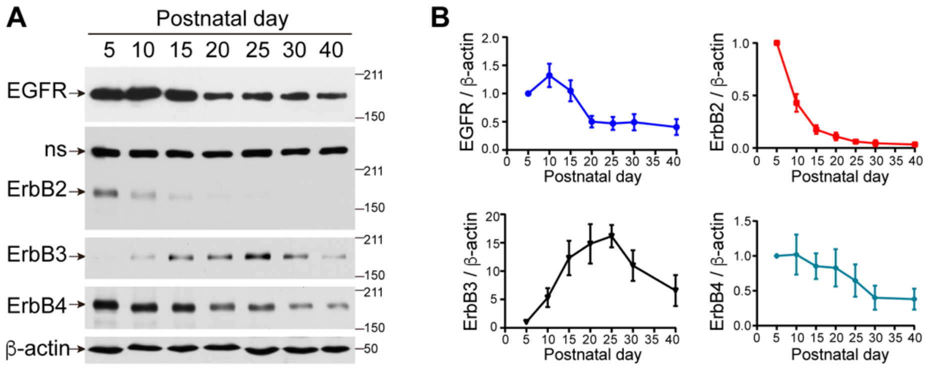


**Supplementary Figure 1: Expression of ErbB receptor members in white matter regions at different postnatal days. A** Western blot results of ErbB receptors in white matter regions isolated from mice at different postnatal days. ns, non-specific band. **B** Quantitative data of the western blot results in **A**. *n* = 3 independent experiments for each postnatal day.

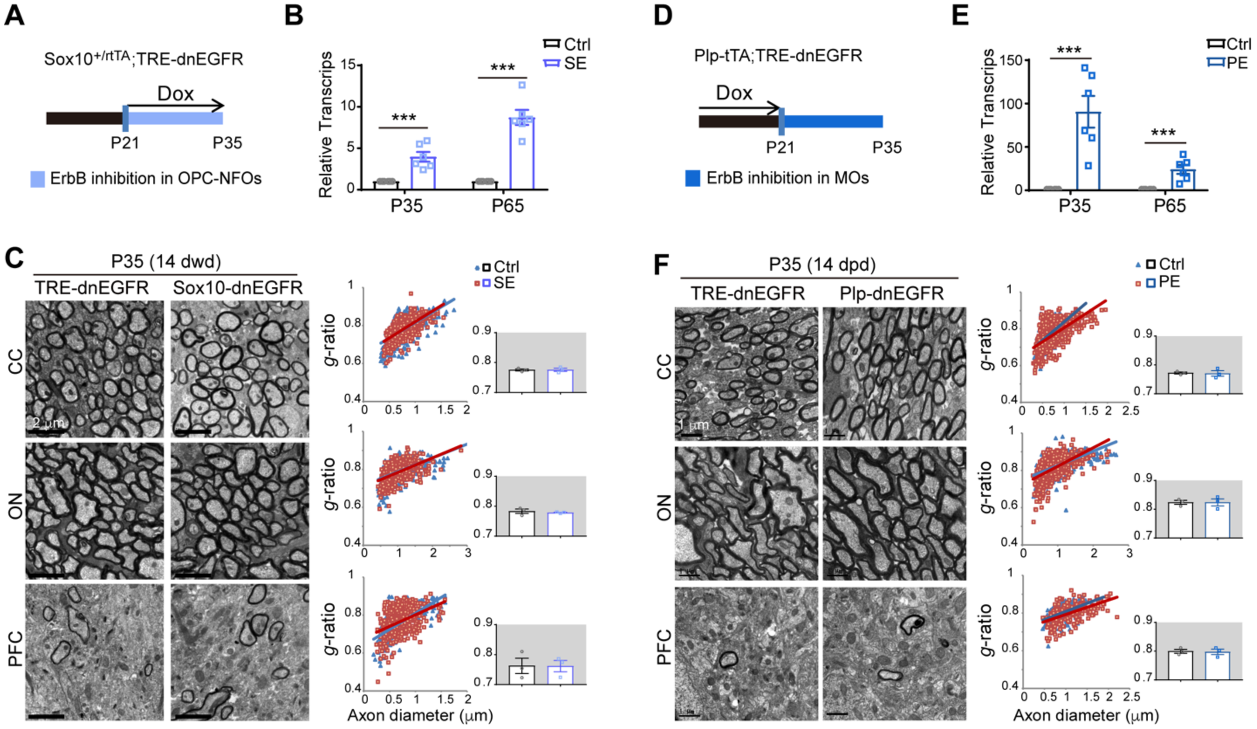


**Supplementary Figure 2: No myelin alteration in *Sox10*-dnEGFR (SE) or *Plp*-dnEGFR (PE) mice at P35 with 14 days of Dox treatment. A**, **D** Dox treatment setting for indicated mice and littermate controls. **B**, **E** Without an antibody recognizing dnEGFR specifically, the expression of dnEGFR/EGFR was examined by real-time RT-PCR. As *Sox10*$^{+/\mathrm{rtTA}}$ targets a transient cellular stage, transcripts of dnEGFR/EGFR in the subcortical white matter of *Sox10*-dnEGFR mice were only 4- and 9-fold more than that of the endogenous EGFR in littermate controls (Ctrl) at P35 and P65, respectively (**B**), while that of *Plp*-dnEGFR mice were 90- and 24-fold more than the endogenous EGFR transcripts in littermate controls at P35 and P65, respectively (**E**). *n* = 6 for each group. **C**, **F** EM images of the corpus callosum (CC), optic nerve (ON), and prefrontal cortex (PFC) from *Sox10*-dnEGFR and littermate controls at P35 with 14 dwd (**C**), or from *Plp*-dnEGFR and littermate controls at P35 with 14 dpd (**F**). *g*-ratio was calculated for myelinated axons. *n* = 3 for each group.

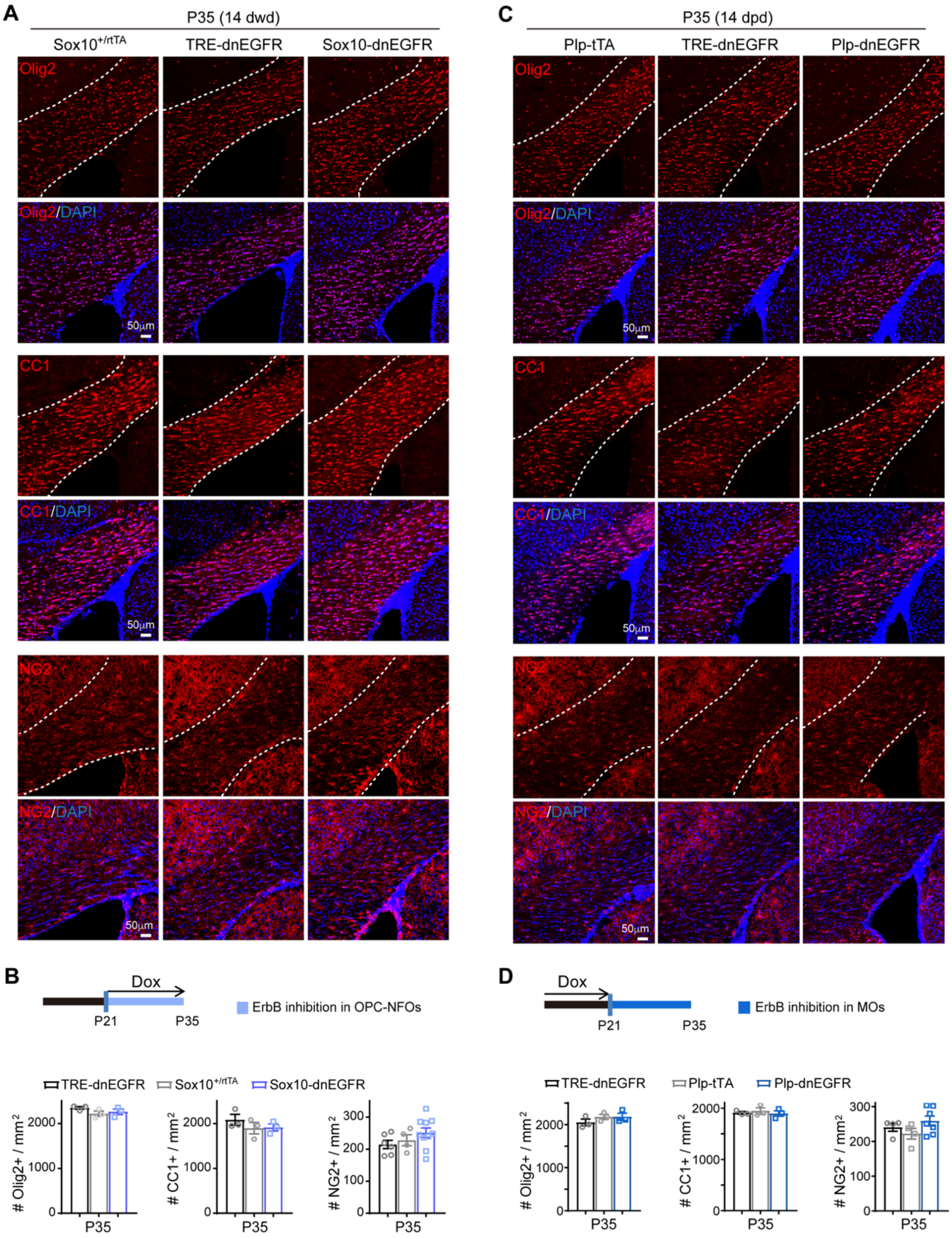


**Supplementary Figure 3: No OL number changes in *Sox10*-dnEGFR or *Plp*-dnEGFR mice at P35 with 14 days of Dox treatment. A**, **C** Immunostaining results of Olig2$^+$, CC1$^+$, and NG2$^+$ cells in the corpus callosum of indicated mice at P35. **B**, **D** Statistic results of Olig2$^+$, CC1$^+$, and NG2$^+$ cell densities in the corpus callosum of indicated mice. Data were analyzed by one-way ANOVA test. For Olig2 and CC1, *n* = 3 mice for each group. For NG2, data were from repeated staining of 3 mice for each group.

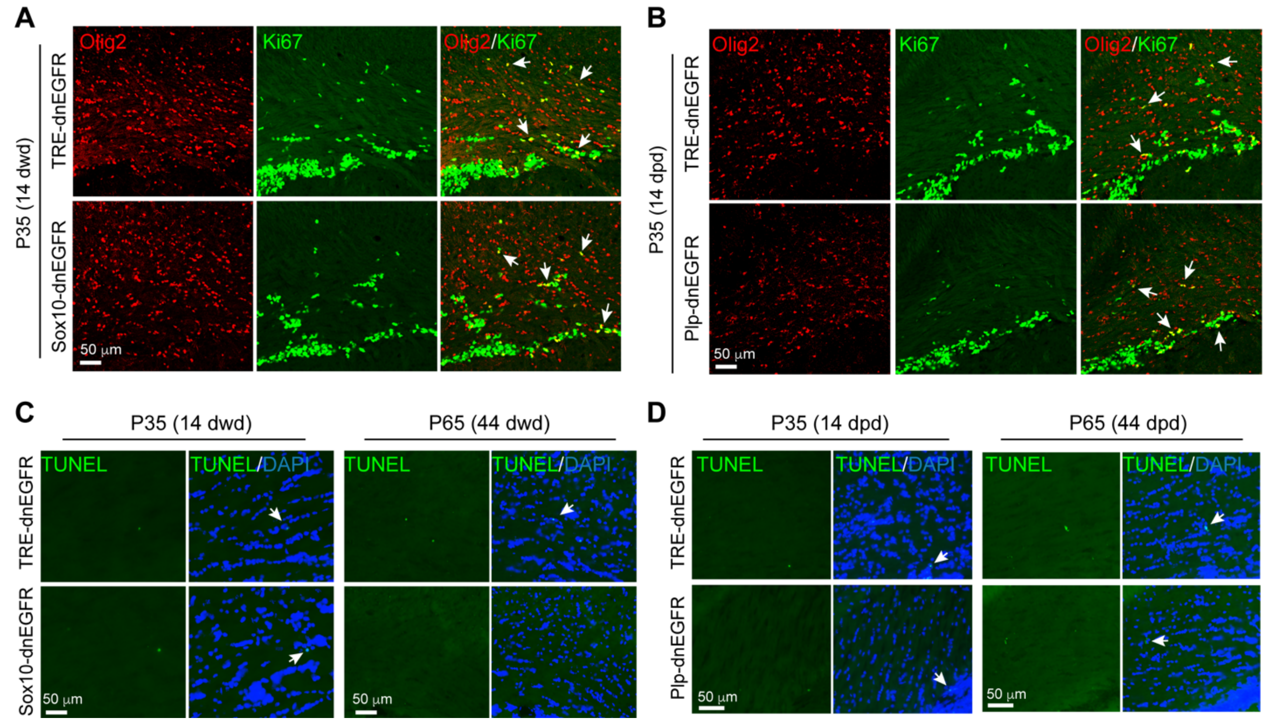


**Supplementary Figure 4: OPC proliferation is not altered in *Sox10*-dnEGFR mice at P35 with 14 days of Dox treatment. A**, **B** Double immunostaining results of Olig2 and Ki67 in the corpus callosum of indicated mice. Arrows, representative double positive nuclei. **C**, **D** Apoptotic cells (TUNEL$^{+}$, white arrows) in the corpus callosum of *Sox10*-dnEGFR mice (**C**), or *Plp*-dnEGFR mice (**D**), were as few as that in littermate controls at indicated ages.

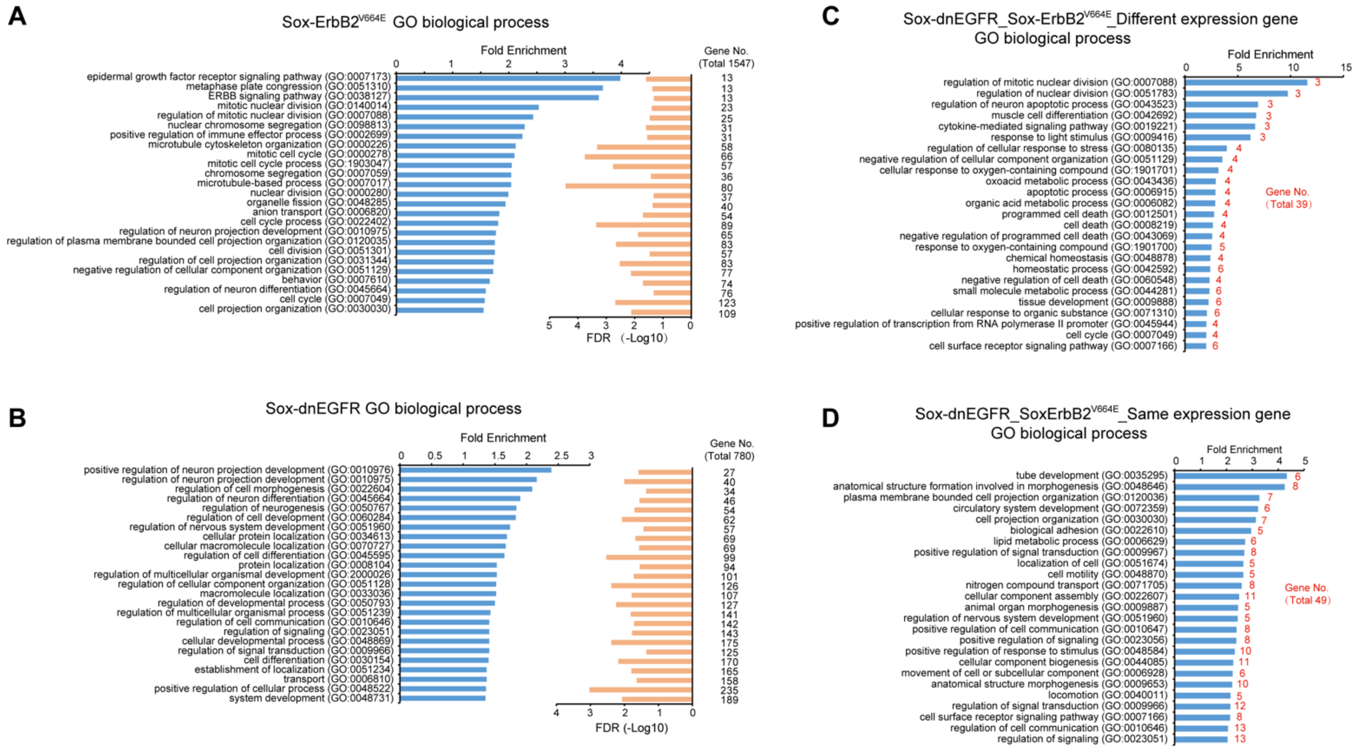


**Supplementary Figure 5: RNA-seq analysis of *Sox10*-ErbB2$^{V664E}$ or *Sox10*-dnEGFR mice with their littermate controls**. **A**, **B** GO enrichment analysis of genes with significant differences in expression between the white matter tissues isolated from *Sox10*-ErbB2$^{V664E}$ mice and that from littermate controls (**A**), or those between *Sox10*-dnEGFR mice and littermate controls (**B**). GO terms with enrichment fold at top 25 were plotted as blue bar at the left. -Log10 of FDR value by Fisher's Exact Test of the same term was plotted as orange bar at the right. Gene numbers within each term were illustrated at the right side. **C**, **D** GO enrichment analysis of genes with different expression tendencies (**C**) or same expression tendencies (**D**) in *Sox10*-dnEGFR and *Sox10*-ErbB2$^{V664E}$ mice. GO terms with enrichment fold more than 2 and gene numbers at top 25 were plotted as blue bar. Gene numbers within each term were illustrated in red.

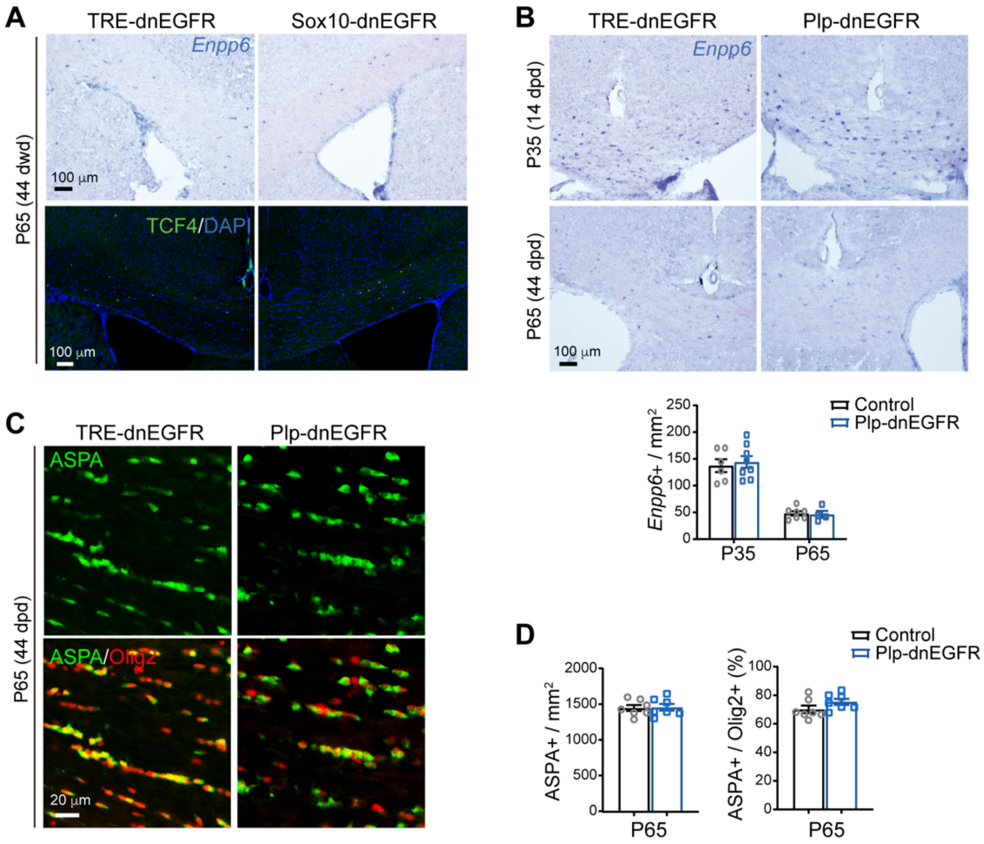


**Supplementary Figure 6: ErbB inhibition does not alter the NFO and MO numbers of *Plp*-dnEGFR mice. A** In situ hybridization results of *Enpp6* and immunostaining results of TCF4 in the corpus callosum of *Sox10*-EGFR mice and littermate controls at P65 with 44 dwd. **B** In situ hybridization results of *Enpp6* in the corpus callosum of *Plp*-dnEGFR mice and littermate controls at indicated ages. Repeated staining results from 3 mice for each group were statistically analyzed. **C** Immunostaining results of MOs ($ASPA^+Olig2^+$) in the corpus callosum of *Plp*-dnEGFR with their littermate controls at indicated ages. **D** Statistic results of $ASPA^+$ cell densities and the ratio of $ASPA^+$ to $Olig2^+$ cell numbers in the corpus callosum of *Plp*-dnEGFR mice. Data were from repeated staining results from 3 mice for each group.

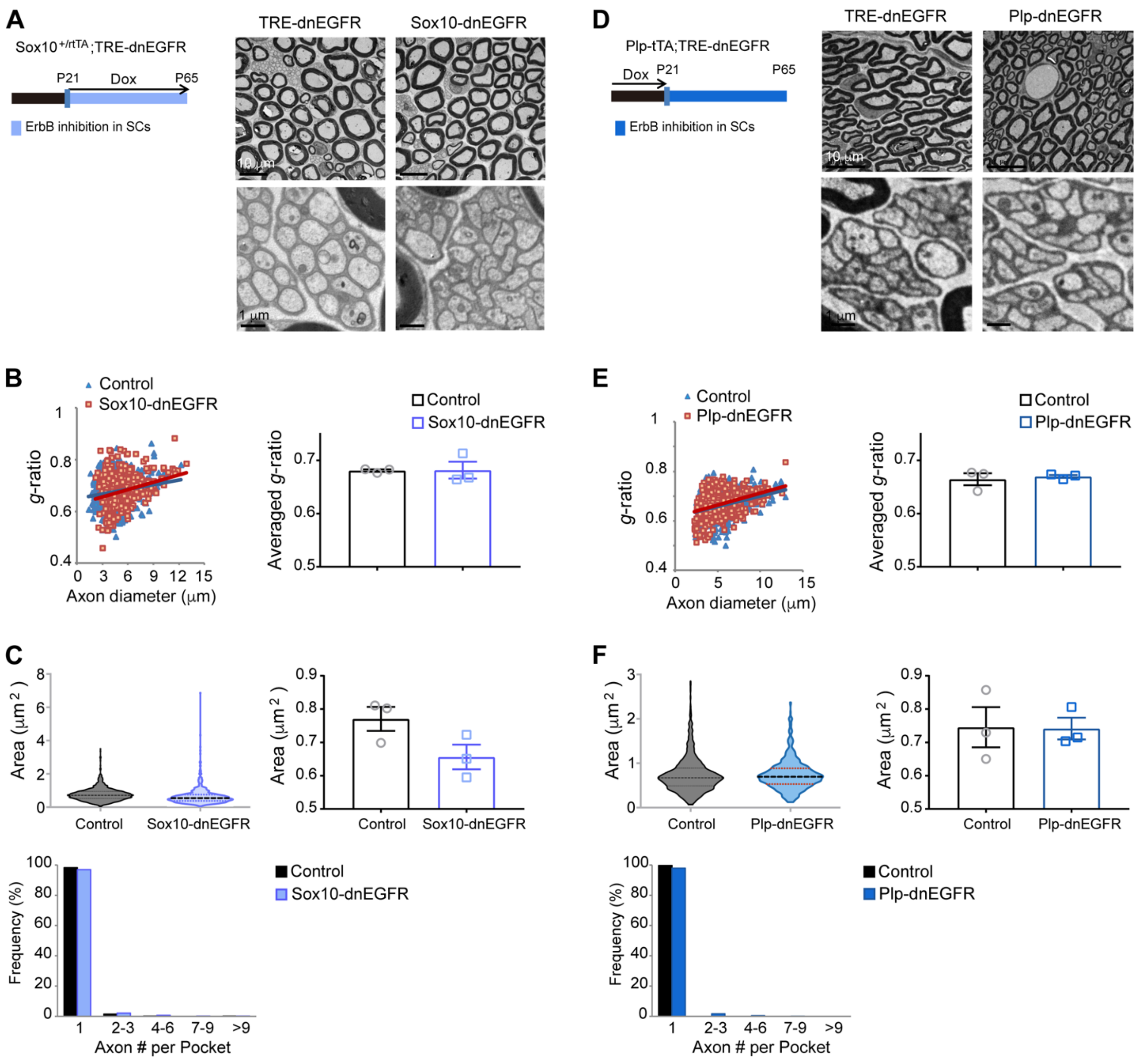


**Supplementary Figure 7: No significant deficiency in peripheral nerves of *Sox10*-dnEGFR or *Plp*-dnEGFR mice after Dox treatment**. Myelin thickness and *g*-raio of myelinated axons, or axonal size and ensheathment of unmyelinated axons, in sciatic nerves were examined by EM and analyzed for *Sox10*-dnEGFR with littermate controls (**A**-**C**), or *Plp*-dnEGFR with littermate controls (**D**-**F**). **A**, **D** The top layers of EM images showed myelinated axons in the sciatic nerves, while the bottom layers of EM images showed unmyelinated axons. **B**, **E** *g*-ratio of myelinated axons were analyzed. $n$ = 3 for each group. **C**, **F** Top left: Axonal sizes of unmyelinated axons were measured by their areas in cross sections, and data were shown as violin plots. Top right: averaged areas of unmyelinated axons of each mouse. Bottom: For the ensheathment analysis of unmyelinated axons, axon numbers in each pocket were counted and quantified by frequency. For mature peripheral nerves, the majority of non-myelinating SC pockets only ensheathe one axon. Data were from 3 mice for each group.

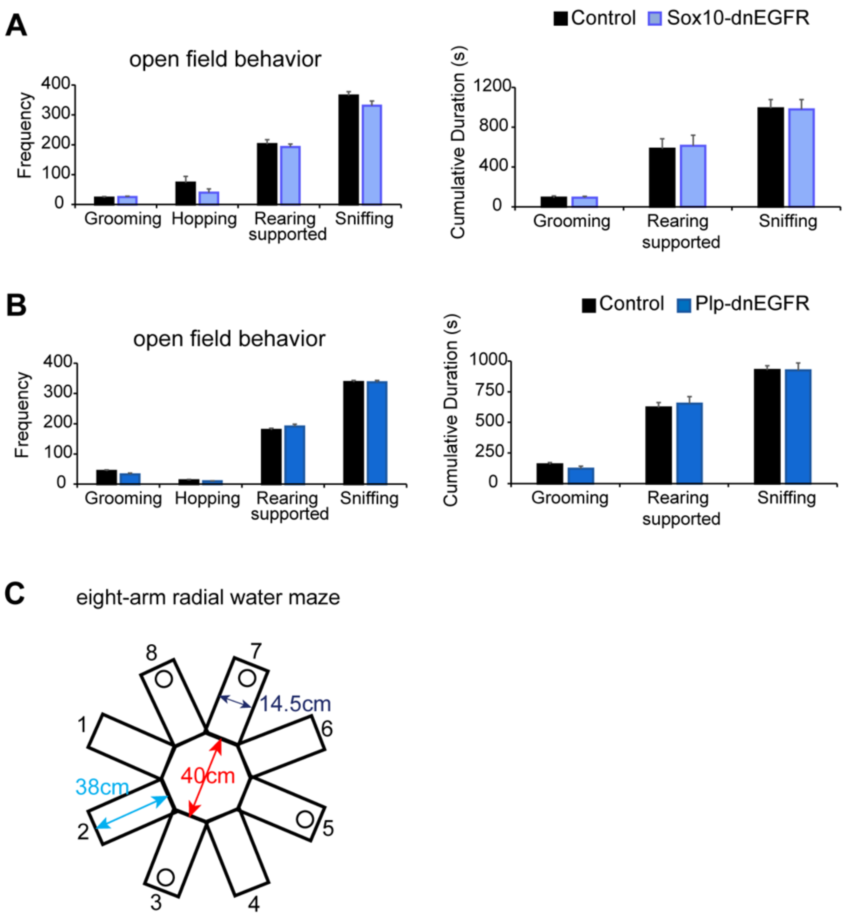


**Supplementary Figure 8: No stereotyped behavioral abnormalities revealed for *Sox10*-dnEGFR or *Plp*-dnEGFR mice. A**, **B** Stereotyped behaviors of adult *Sox10*-dnEGFR mice with littermate controls (**A**), or *Plp*-dnEGFR mice with littermate controls (**B**), were analyzed in the open field test and exhibited no difference. control $n = 11$, *Sox10*-dnEGFR $n = 13$ (**A**); control $n = 19$, *Plp*-dnEGFR $n = 14$ (**B**). Data were analyzed by multiple $t$ test without correction for multiple comparisons. **C** Illustration showing the setting for eight-arm radial water maze. Four hidden platforms were placed at the end of a same set of arms with 38-cm distance to the central zone at the training and test days. Mice started swimming with face to the arm end from No.1 arm in each trial, and the visited platform was removed before the next trial after 30-sec gap.

### Other Supplementary Files:

**Supplementary Video 1:** Performances recorded for *Sox10*-dnEGFR mice and *Plp*-dnEGFR mice, as well as their controls, in the 4th trial of eight arm radial water maze at the test day.

**Supplementary Table 1:** Processed RNA-seq results of genes with differential expression in white matter tissues between *Sox10*-ErbB2$^{V664E}$ mice and littermate *TRE*-ErbB2$^{V664E}$ mice at P30 with 9 dwd.

**Supplementary Table 2:** Processed RNA-seq results of genes with differential expression in white matter tissues between *Sox10*-dnEGFR mice and littermate *TRE*-dnEGFR mice at P35 with 14 dwd.
